# Layer-specific integration of locomotion and concurrent wall touching in mouse barrel cortex

**DOI:** 10.1101/265165

**Authors:** Asli Ayaz, Andreas Stäuble, Aman B Saleem, Fritjof Helmchen

## Abstract

During navigation rodents continually sample the environment with their whiskers. How locomotion modulates neuronal activity in somatosensory cortex and how self-motion is integrated with whisker touch remains unclear. Here, we used calcium imaging in mice running in a tactile virtual reality to investigate modulation of neurons in layer 2/3 (L2/3) and L5 of barrel cortex. About a third of neurons in both layers increased activity during running and concomitant whisking, in the absence of touch. Fewer neurons were modulated by whisking alone (<10%). Whereas L5 neurons responded transiently to wall-touching during running, L2/3 neurons showed sustained activity after touch onset. Consistently, neurons encoding running-with-touch were more abundant in L2/3 compared to L5. Few neurons across layers were also sensitive to abrupt perturbations of tactile flow. We propose that L5 neurons mainly report changes in touch conditions whereas L2/3 neurons continually monitor ongoing tactile stimuli during running.

## INTRODUCTION

We sense the outside world through the continuous interactions between the streams of sensory inputs and motor actions. How sensory and motor information are integrated in the brain is critical for understanding sensory processing during behavior. Recent studies in head-restrained mobile mice explored effects of locomotion on sensory processing and found increased spontaneous and evoked activity of excitatory neurons in both primary visual cortex (V1) (Ayaz et al., 2013; Niell and Stryker, 2010) and the lateral geniculate nucleus of the thalamus (Erisken et al., 2014). These effects cannot be explained by simple gain modulation or additive factors because the modulation is often non-monotonic (Ayaz et al., 2013; Erisken et al., 2014; Saleem et al., 2013) and effects are heterogeneous for distinct GABAergic interneuron subtypes (Fu et al., 2014; Pakan et al., 2016; Polack et al., 2013). Further, it is unclear the extent to which locomotion-related increases are specific to the visual system, or generalize to other sensory cortices. In fact, locomotion suppresses excitatory neurons in auditory cortex (Schneider et al., 2014). The effects of locomotion on somatosensory processing have so far been investigated only incidentally (Fu et al., 2014; Sofroniew et al., 2015). Rather than running, whisking behavior has been the main focus of studies on sensorimotor integration in vibrissal primary sensory cortex (S1 or ‘barrel cortex’) (Crochet and Petersen, 2006; Deschênes et al., 2012; Eggermann et al., 2014; Gentet et al., 2010; Petersen, 2014; Poulet and Petersen, 2008; Poulet et al., 2012; Sofroniew and Svoboda, 2015). Membrane potential fluctuations of nearby pyramidal neurons were reported to desynchronize during active whisking without significant changes in firing rate (Crochet and Petersen, 2006; Gentet et al., 2010; Poulet and Petersen, 2008). In addition, whisking was found to modulate different interneuron classes distinctly (Gentet et al., 2010, 2012) and to increase thalamic activity (Urbain et al., 2015).

In their natural environment rodents utilize their whiskers during navigation. The occurrence of running and whisking are typically highly correlated (Arkley et al., 2014; Mitchinson et al., 2011; Sofroniew et al., 2014; Wineski, 1983). Somatosensory processing is likely to vary between running and resting state because behavioral requirements are different: During locomotion mice need to monitor their location, while remaining sensitive to sudden changes in the environment, such as encountering an obstacle on their way. However, while being stationary and exploring an object in detail, mice may emphasize more subtle aspects of somatosensation and screen surface texture or shape. How locomotion state and touch-evoked neuronal responses are integrated in S1 neural circuitry remains an open question.

Integration of visual stimulation and running has been studied in neurons located in either superficial (supragranular) layers (Fu et al., 2014; Keller et al., 2012; Leinweber et al., 2017; Niell and Stryker, 2010) or in deeper (infragranular) layers (Ayaz et al., 2013; Saleem et al., 2013). Different layers of sensory cortices harbor distinct input and output patterns. For example, afferent inputs to barrel cortex from the ventral posterior medial (VPM) nucleus in thalamus mainly target L4 (plus L5/6; c.f.Constantinople and Bruno, 2013), whereas axons from the posterior medial (POM) nucleus terminate in L1 and L5A (Lu and Lin, 1993; Petreanu et al., 2009; Wimmer et al., 2010). In addition, long-range projections between S1 and other cortical areas, e.g., vibrissal motor cortex (M1) and secondary somatosensory cortex (S2), show significant layer specificity (Aronoff et al., 2010; Kinnischtzke et al., 2014; Mao et al., 2011; Petreanu et al., 2009). Such laminar-specific connectivity suggests different functional roles of superficial and deep-layer neurons in S1. In line with this notion, recent studies that directly compared sensory processing across layers have started to uncover functional distinctions within the laminar organization (van der Bourg et al., 2016; de Kock and Sakmann, 2009; O’Connor et al., 2010; Pluta et al., 2015; Sofroniew et al., 2015).

## RESULTS

### Calcium imaging across barrel cortex layers in a tactile virtual reality

To study locomotion effects on touch processing in barrel cortex, we built a tactile virtual reality setup to mimic running and whisking along a wall in the dark (Figure 1A; Experimental Procedures). Mice were head-fixed on top of a ladder wheel under a two-photon microscope. We continually recorded run speed and induced touches by bringing a sandpaper-covered rotating cylinder (‘textured-wall’) in contact with the whiskers on one side of the face. Whisker movements were imaged with a high-speed camera. This setup enabled us to consider several experimental conditions (Figure 1B): First, the mouse was free to run or rest on the treadmill in the absence of the textured-wall (‘No-touch’). Second, the texture was moved in contact with the whiskers after the mouse had run a predefined distance, rotating at the same speed as the animal’s run speed (‘Closed-loop’). Third, texture speed and run speed were decoupled (‘Open-loop’). The varying combinations of texture and run speed in this ‘Open-loop’ condition allowed us to dissect their contributions to neuronal activity in S1. Finally, to explore neuronal responses to an abrupt mismatch of tactile flow and run speed, we applied brief perturbations during Closed-loop trials by halting the rotating texture for 2 seconds at a random time point during the touch. In all conditions, mice were free to move on the treadmill and did not receive any reward. To avoid whisker stimulation by the treadmill, the bottom rows of whiskers were trimmed. Measurements were made in several experimental sessions spread over 3 weeks (see Experimental Procedures).

**Figure 1.**
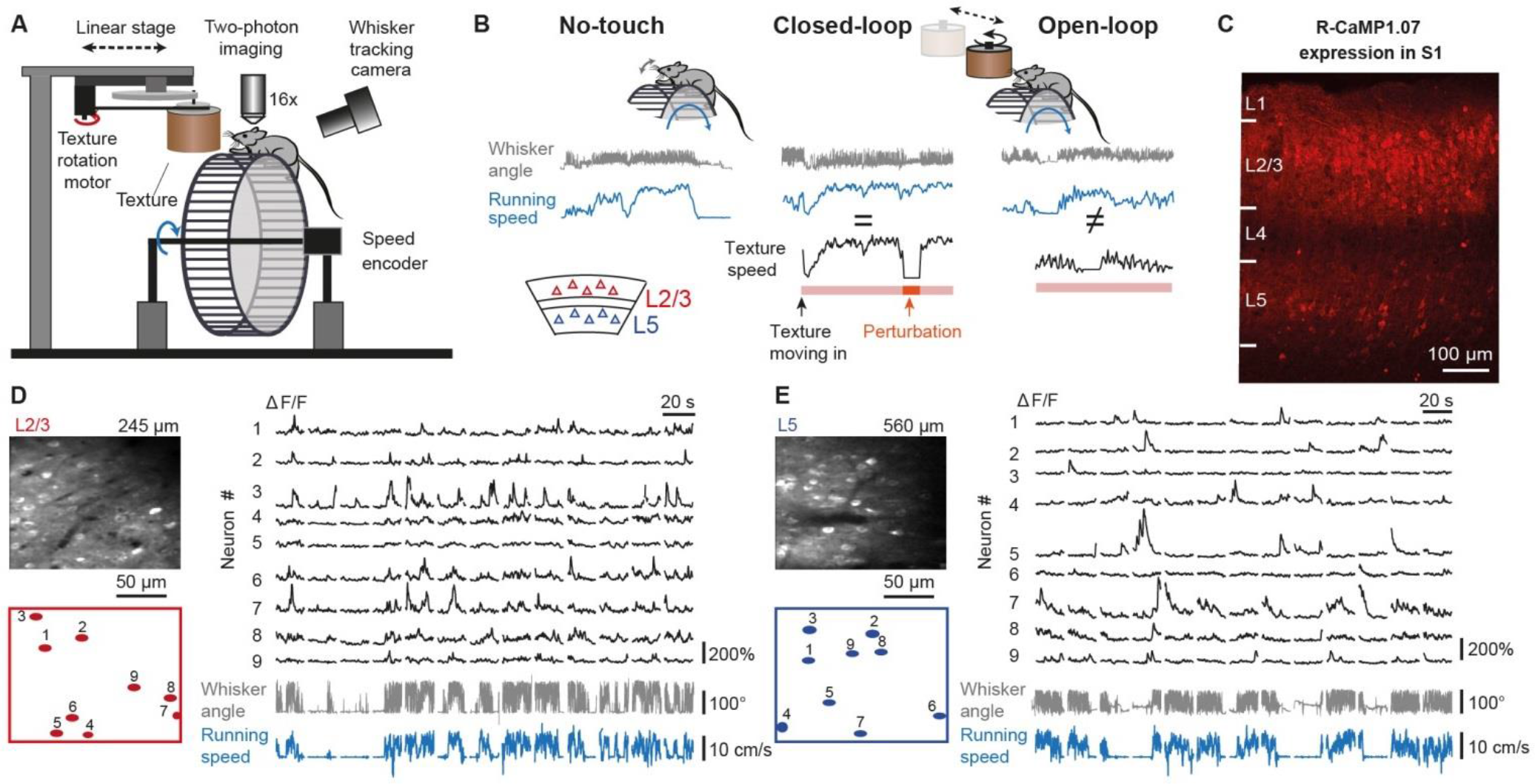
Calcium imaging in L2/3 and L5 of mouse barrel cortex during various running and whisking conditions. (A) Schematic of virtual reality setup with a head-restrained mouse on top of a rung-ladder treadmill. A sandpaper-covered cylinder (‘texture’) can be moved in contact with the whiskers. Run speed is tracked with an encoder and the mean whisker angle is monitored with a high-speed video camera. (B) Example traces of whisker angle (gray), run speed (blue), and texture rotation speed (black), illustrating the 3 experimental conditions of ‘No-touch’, ‘Closed-loop’, and ‘Open-loop’. The pink bottom bar indicates when the texture contacts the whiskers. The orange segment highlights the intermittent halt of texture rotation to introduce a brief perturbation period in Closed-loop trials by uncoupling run speed and texture speed. (C) Confocal image of virally-induced R-CaMP1.07 expression pattern in a coronal slice of somatosensory cortex of a wild type mouse. (D) Left: In vivo two-photon image of R-CaMP1.07-expressing L2/3 neurons with selected ROIs below. Right: ΔF/F calcium transients of 9 example neurons with simultaneously recorded mean whisker angle (gray) and running speed (blue) below. (E) Same as in D but for example L5 neurons in SI.

To measure neuronal activity across cortical layers we injected AAV2.1-EFα1-R-CaMP1.07 into barrel cortex of adult mice, causing expression of the genetically encoded calcium indicator R-CaMP1.07 (Bethge et al., 2017; Ohkura et al., 2012) in L2/3 and L5 neurons (Figure 1C; Experimental Procedures). R-CaMP1.07-expressing cells presumably mainly represent pyramidal neurons as similar viral vectors previously barely drove expression in GABAergic interneurons (Chen et al., 2013). Using the lack of expression in L4 we selected imaging areas either above or below this laminar landmark (6 areas each in L2/3 and L5; imaging depths: 221-385 µm for L2/3 and 450-664 µm for L5; n = 5 mice). In total, we measured calcium signals in 426 neurons in L2/3 and 275 neurons in L5. Figure 1D and E show example calcium transients measured in neuronal subpopulations in L2/3 and L5 in freely running mice. Assuming single action potential-evoked R-CaMP1.07 ΔF/F changes of ~7% amplitude with ~0.3 s decay time constant (Bethge et al., 2017) and linear summation, a 50% steady-state ΔF/F change reflects a firing rate change of about 24 Hz (Helmchen et al., 1996) whereas sharp large calcium transients indicate the occurrence of bursts with variable numbers of action potentials.

Here, we investigated whether L2/3 and L5 neurons of barrel cortex are modulated during running and whether they integrate sensory and motor signals differently. We applied two-photon calcium imaging in head-restrained mice running along a “virtual tactile wall”. This setting allowed us to study the integration of two motor behaviors—‘whisking’ and ‘running’—with prolonged sensory touches. We find that running behavior strongly modulates neuronal activity and touch-evoked responses in barrel cortex and that this modulation cannot be explained by the accompanying whisking behavior. Furthermore, we reveal distinct features of L2/3 and L5 neurons regarding the representation of prolonged touch stimuli and the integration of self-motion and touch.

### Running increases barrel cortex activity in the absence of a wall

We first investigated how locomotion and whisking affect the activity of S1 neurons in the absence of texture touch (No-touch sessions). Based on the recorded running speed and measured whisker movements, we defined four different behavioral states (Figure 2A; Experimental Procedures): Animals spent about half of their time running, which was always accompanied by simultaneous whisking as reported previously (Sofroniew et al., 2014). Stationary periods included whisking as well as no-whisking episodes whereas animals almost never ran without whisking. To assess activity changes induced solely by whisking we excluded all running periods and compared the mean fluorescence change during episodes of whisking and no-whisking for each neuron. Whisking barely modulated the mean activity in L2/3 and L5 populations (Figure 2B and Figure S1). To account for differences in activity levels we calculated a whisking modulation index (MI) for each neuron (as defined in Pakan et al., 2016, see Experimental Procedures). Whisking MIs were close to but significantly different from zero (p < 10-4 for L2/3 and L5, Wilcoxon signed rank test) and not different for L2/3 and L5 neurons (Figure 2C; 0.071 ± 0.009, n = 342, and 0.055 ±0.013, n = 168, respectively; mean ± s.e.m.; p > 0.05, Wilcoxon rank sum test). As alternative analysis we averaged neuronal fluorescence changes after alignment to detected onsets of whisking while the animal was stationary (Figure 2D,E; considering only fluorescence trace segments until whisking stopped). Consistent with a weak modulation by whisking, only a minor fraction of neurons (3%, 11/342, in L2/3 and 8%, 14/168, in L5) showed significant fluorescence increases upon whisking onset (Figure 2D,E and Figure S1).

**Figure 2.**
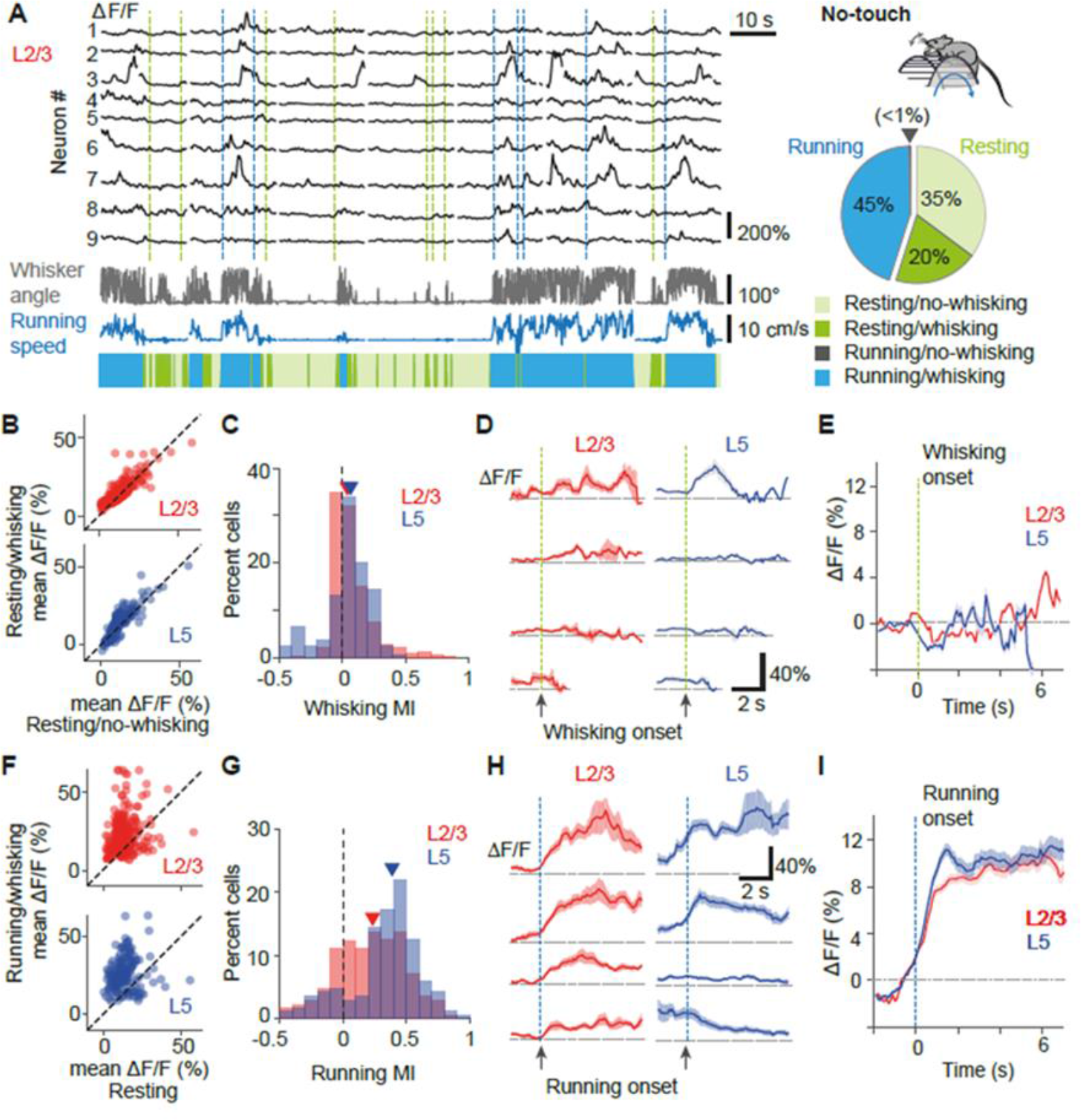
Running with concomitant whisking increases L2/3 and L5 activity more than whisking alone. (A) Left: Example ΔF/F traces along with whisker angle and running speed in the initial No-touch trials of Figure 1D. The 4 possible running and whisking state conditions are color-coded. Green and blue vertical dotted lines mark whisking onset (only during resting periods) and running onset, respectively. Right: Pie chart of the distribution of times spent in the 4 states for 5 mice (only ‘No-touch’ condition). Gray triangle indicates <1% fraction of time spent in the running/no-whisking state. (B) Scatter plots of mean ΔF/F amplitude in resting/whisking periods versus resting/no-whisking periods for 342 L2/3 neurons (red) and 168 L5 neurons (blue). Dashed lines indicate unity lines. (C) Distribution of whisking modulation index (MI) for L2/3 and L5 neurons. Red and blue triangles indicate medians. (D) Average ΔF/F traces of example neurons aligned to whisking onset (green vertical dotted line) during resting periods (shading shows ±s.e.m.). Only the period of continued whisking after onset was considered for each trial, hence the variable lengths of traces for different neurons. (E) Population average of whisking-onset aligned ΔF/F traces for all recorded neurons. (F) Scatter plots of mean ΔF/F amplitude in running/whisking periods versus resting periods for L2/3 (top) and L5 (bottom) neurons. (G), (H), (I), Analogue plots to C, D, and E for running modulation index (MI) and running-onset aligned ΔF/F traces.

Similarly, we examined how running modulates activity of S1 neurons. As animals almost always whisked during locomotion (Figure 2A; cf. Sofroniew et al., 2014) any running period also included whisking (referred to as ‘running/whisking’). For each neuron we compared mean activity during running/whisking and during resting periods (including both ‘whisking’ and ‘no-whisking’ episodes). Running/whisking caused significant mean ΔF/F increases in the majority of both L2/3 and L5 neurons and decreases in some neurons (Figure 2F and Figure S1). Consistently, a running MI—defined in analogy to the whisking MI— revealed increased activity during running/whisking for both populations (Figure 2G; running MI 0.210 ± 0.016, n = 342, for L2/3 and 0.329 ± 0.023, n = 168, for L5 neurons, respectively; p < 10-4 for both populations, Wilcoxon signed rank test) with significantly larger running modulation for L5 neurons (p < 10-4, Wilcoxon rank sum test). We also computed average fluorescence changes aligned to locomotion onsets, considering only fluorescence trace segments until running stopped (Figure 2H). About one third of neurons showed a significant increase in activity upon running onset (32%, 108/342, in L2/3 and 38%, 64/168, in L5; Figure S1D). This increase persisted over several seconds as reflected in the population average of run-onset aligned responses of L2/3 and L5 neurons (Figure 2I). Taken together, the No-touch sessions revealed that running/whisking increases the activity of both L2/3 and L5 neurons in barrel cortex to a larger extent than whisking alone

### Sustained versus transient responses to continuous wall touch in L2/3 and L5 neurons

After determining the effect of running and whisking in the absence of sensory stimulation, we next compared responses evoked by texture touch in the Closed-loop condition. In each trial, after the mouse had run a predefined distance, a textured wall moved in contact with the whiskers, rotating at the same speed as the animal was running (Figure 3A). We aligned the fluorescence traces to ‘touch onset’, defined as the time point when the texture started moving towards the whiskers (Experimental Procedures). More than half of the neurons were responsive to wall touch (64%, 268/420, in L2/3 and 57%, 157/275, in L5; see Experimental Procedures). Both L2/3 and L5 neurons displayed a strong initial response to touch. However, L2/3 neurons continued to exhibit a sustained response to ongoing wall touch whereas L5 neurons showed a transient response (Figure 3B,C). Although a rough-textured wall elicited slightly larger activity than a smooth-textured wall both in L2/3 and L5 neurons, their difference in temporal response profile remained the same, independent of texture identity (Figure S2). In an early response window the peak ΔF/F change relative to the mean ΔF/F value in a pre-touch period was 32 ± 0.9% and 26 ± 0.9% for responsive L2/3 and L5 neurons, respectively (mean ± s.e.m.; values were significantly different, p < 10-4, Wilcoxon rank sum test, Figure 3B). In a late response window (2 – 3 s after touch onset) the ΔF/F traces showed a strong decrease in L5 but not in L2/3 (Figure 3C). For quantification we calculated the level of ΔF/F suppression in the late window relative to the value in the early window (Figure 3C). Suppression was significantly larger for L5 compared to L2/3 neurons (69.6 ± 4.4% for L5 and 16.6 ± 2.7% for L2/3, p < 0.0001, Wilcoxon rank sum test; Figure 3D).

**Figure 3.**
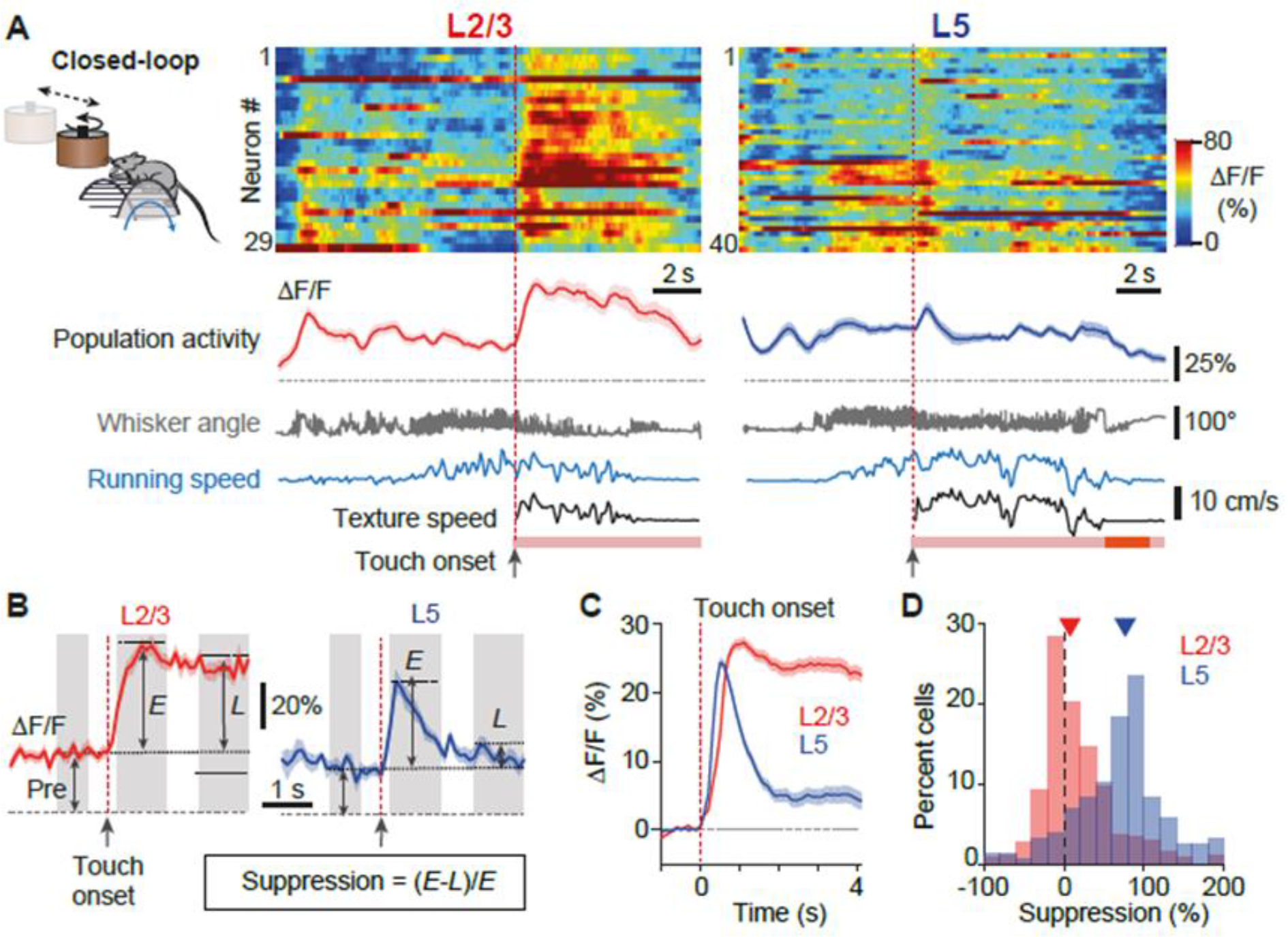
Wall-touching during running evokes sustained responses in L2/3 neurons and transient responses in L5 neurons. (A) Population dynamics in L2/3 (left) and L5 (right) neurons for two example trials under Closed-loop condition. The heat map represents ΔF/F calcium transients in pseudo-color code. Below the population mean ΔF/F trace (solid line, ±s.e.m.) whisker angle (gray), running speed (blue), and texture speed (black) are shown. Touch onsets are indicated by red vertical dotted lines; pink bars represent periods of texture contact and dark orange segment is the period when texture rotation is stalled. (B) Mean ΔF/F traces aligned to touch onset (red dotted line) of an example L2/3 and a L5 neuron. Shaded areas indicate windows for analysis (pre-touch, ‘Pre’, -1 to -0.3 s; early, ‘E’, 0.3 to 1.3 s; late, ‘L’, 2 to 3 s). (C) Population averages (±s.e.m.) of touch-aligned ΔF/F traces for all touch-responsive neurons in L2/3 (red, n = 268/420) and L5 (blue, n = 157/275). Neurons were considered touch-responsive when E > 2σ, where σ is the minimum standard deviation (see Experimental Procedures). (D) Distribution of percent suppression as defined in B for L2/3 and L5 touch-responsive neurons. Arrowheads mark medians of 76.6% and 7.7%, respectively.

In the Closed-loop condition the wall touch almost always happened during running as mice had to travel a certain distance before reaching a wall. To test whether laminar specificity of wall touch responses depends on the locomotion state we also examined touch responses in Open-loop condition (Figure 4A). This allowed us to sort wall touch events into two groups according to whether the touch occurred during running or during resting (see Experimental Procedures). Although neuronal activity overall was lower during resting, in either condition L2/3 neurons gave a sustained response after wall touch whereas L5 neuron activity decreased over time (Figure 4B-D). In summary our findings reveal that touch-evoked activity in barrel cortex shows laminar specificity, with L2/3 neuronal subsets showing sustained activity during continuous touch whereas subsets of L5 neurons respond transiently.

**Figure 4.**
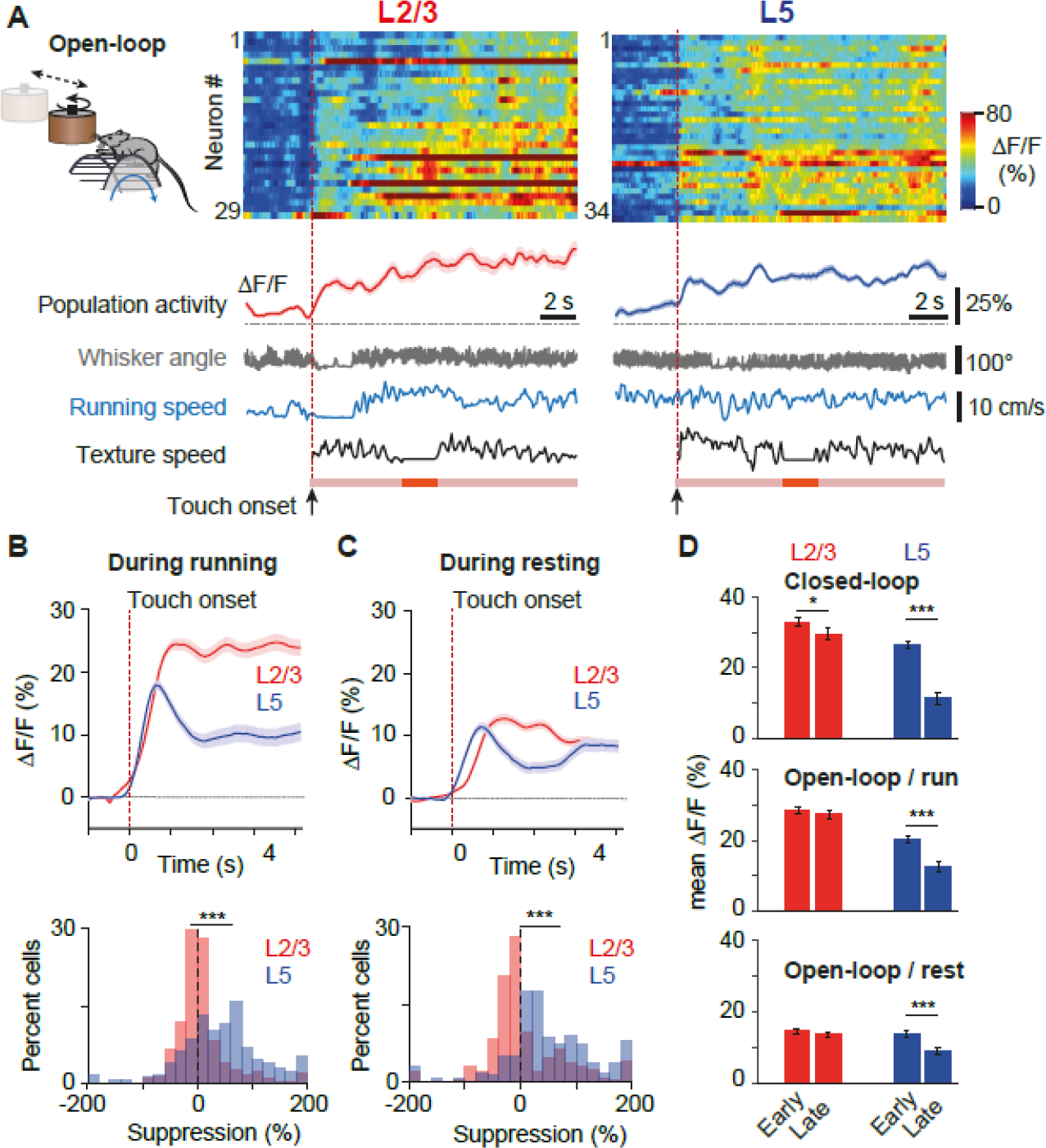
Touch-evoked responses under Open-loop condition. (A) Population dynamics in L2/3 (left) and L5 (right) neurons for two example trials under Open-loop condition. The heat map represents ΔF/F calcium transients in pseudo-color code. Below the population mean ΔF/F trace (solid line, ±s.e.m.) whisker angle (gray), running speed (blue), and texture speed (black) are shown. Touch onsets are indicated by red vertical dotted lines; pink bars represent periods of texture contact; orange segment is the period where texture-rotation is briefly stopped. Imaging field of views are the same as in Figure 3A, with identical neuron numbers for L2/3. In L5 some neurons were not captured in the corresponding imaging session, hence the smaller number of neurons. (B) Of the touch-responsive neurons in Open-loop condition 255 out of 342 of L2/3 (red) and 177 out of 236 L5 (blue) neurons had touch events during running. Top panel: population averages (±s.e.m.) of touch-aligned ΔF/F traces during running. Bottom panel: distribution of percent suppression of touch responses during running. Mean suppression was 8.1 ± 2.9% and 44.1 ± 5.6% for L2/3 and L5 neurons, respectively (p < 0.001, Wilcoxon rank sum test). (C) Same as in B but for touch-responses that occurred during resting (245/342 L2/3 neurons, red, and 167/236 L5 neurons, blue). Mean suppression was 6.4 ± 4.4% for L2/3 and 49.8 ± 6.3% for L5 neurons, respectively (p < 0.001). (D) Comparison of early and late phase touch responses for L2/3 (red) and L5 (blue) populations under 3 conditions: Closed-loop (top panel), Open-loop during running (middle panel) and Open-loop during resting (bottom panel; paired T-test, * p < 0.01, *** p < 0.0001).

### A fraction of neurons is sensitive to mismatch of running speed and tactile flow

In visual cortex a subset of neurons reports mismatches between the animal’s motion and visual flow by increased activity (Keller et al., 2012; Leinweber et al., 2017). Here, we explored whether mismatch-sensitive neurons also exist in barrel cortex. In Closed-loop condition, we introduced strong mismatches between locomotion and tactile flow by abruptly halting the rotating texture for 2 seconds at a random time point during the touch. We found small numbers of neurons that either increased (‘up-modulated’) or decreased (‘down-modulated’) their activity upon such perturbation (example cells shown in Figure 5A,B). For quantification we averaged ΔF/F traces after alignment to the perturbation onset and defined the perturbation response amplitude as the mean ΔF/F change in a 2-s time window after perturbation relative to pre-perturbation baseline (Figure 5B). The distribution of perturbation response amplitudes was similar for L2/3 and L5 neurons, albeit with a slightly but significantly lower mean value for L2/3 neurons (Figure 5C; -4.2 ± 0.6%, n = 420, and -1.7 ± 0.7%, n = 275, for L2/3 and L5, respectively; p < 0.001, Wilcoxon rank sum test). Overall, 4.5% (19/420) of L2/3 and 3.3% (9/275) of L5 neurons showed a significant increase in response to mismatch between running speed and tactile flow (see Experimental Procedures). Significantly down-modulated neurons were more abundant in L2/3 (12.4%, 52/420) than in L5 (6.6%, 18/275) (Figure 5D). Decreased activity was the dominating effect at the population level both in L2/3 and L5 (Figure S3). Whereas decreased activity may be explained by reduced stimulation of the whiskers when the rotation of the texture cylinder is stopped, the increased activity upon texture halt can be considered as representing a true mismatch signal. Our Open-loop experiments also revealed that increased mismatch responses were not a pure sensory response as neurons did not show increased activity to texture stall when animals were at rest whereas the number of mismatch responsive neurons during running was similar to Closed-loop condition (Figure S4).

**Figure 5.**
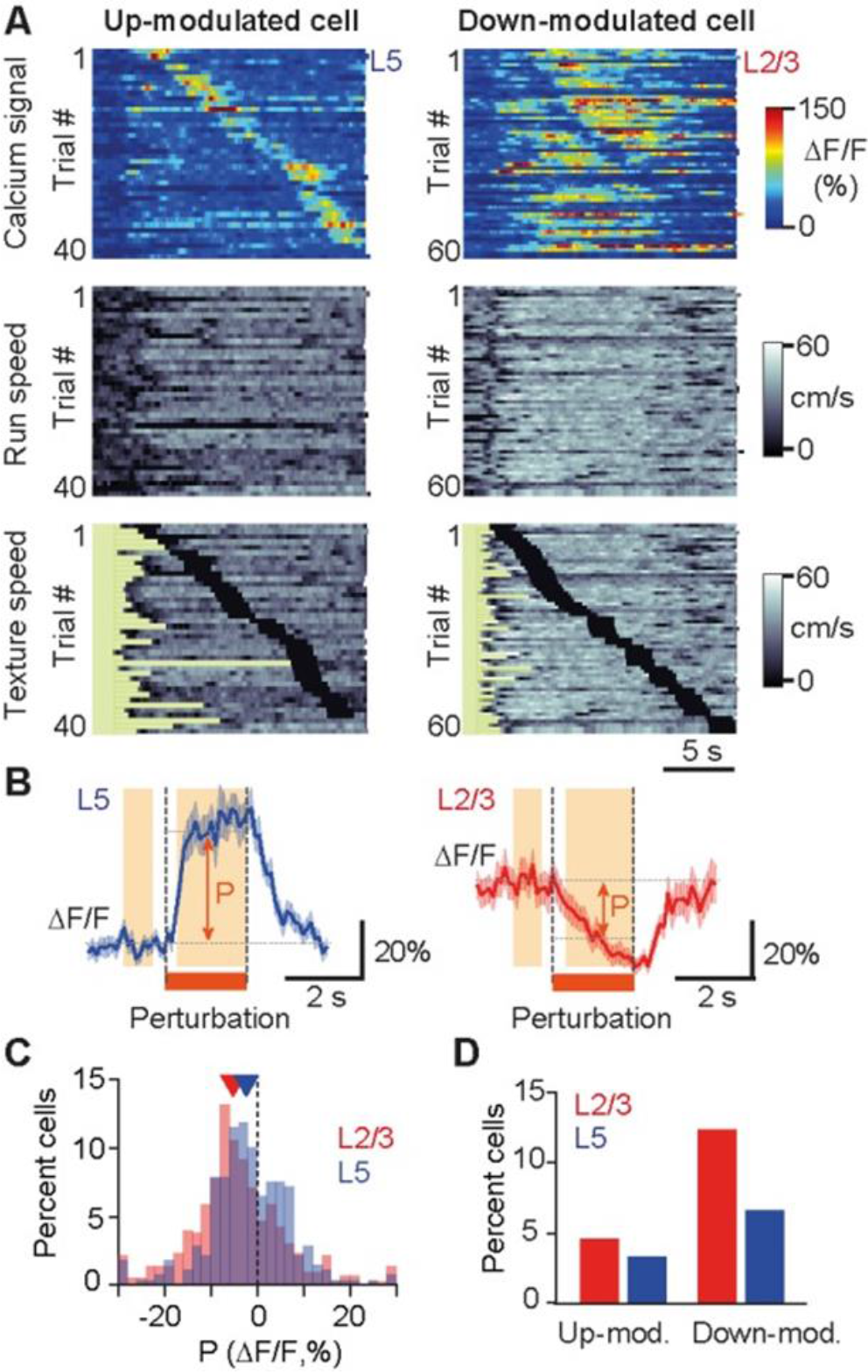
A subset of neurons responds to perturbations of tactile flow. (A) Calcium signals for two example neurons across multiple Closed-loop trials. Trials are sorted according to when the perturbation (2-s texture halt) occurred. Running speed and texture speed are plotted below with black periods marking stop of texture rotation. The left L5 neuron is an example of an up-modulated cell whereas the right L2/3 neuron exemplifies down-modulation upon perturbation. (B) Mean ΔF/F traces (± s.e.m.) aligned to perturbation onset for the example neurons in A. ‘P’ indicates the amplitude of perturbation-induced modulation defined as the difference between mean ΔF/F values before (−1 to -0.3 s window) and during perturbation (+0.3 to 2 s, orange shaded areas). (C) Distribution of perturbation-induced modulation for all L2/3 (red, n = 420) and L5 (blue, n = 275) neurons. Triangles mark medians of -5.1% and -2.3% for L2/3 and L5 neurons, respectively. (D) Percentage of significantly up-modulated (P > 2σ) and down-modulated (P < -2σ) neurons in L2/3 and L5 populations, respectively, with σ denoting baseline noise (Experimental Procedures).

### Neurons integrating self-motion and touch are more numerous in L2/3 than in L5

After revealing neuronal population responses to salient events (e.g., running onset, touch, and perturbation) we evaluated how homogenously these sensorimotor aspects are functionally represented among L2/3 and L5 neurons in barrel cortex. To this end, we considered Open-loop experiments during which various combinations of running and touch occurred. Some neurons faithfully increased activity when the animal touched the wall, independent of running (‘Touch cells’; Figure 6A). Another subset of neurons was most active when the animal was running independent of the wall touch (‘Run cells’). A relatively large population of neurons was most responsive when wall touch and running occurred concomitantly (‘Integrative cells’). Finally, some rare neurons were most active when the animal was stationary in the absence of wall touch (‘Rest cells’). To classify neurons according to these categories, we computed the mean ΔF/F values for the four combinations of locomotion and touch state (‘no-running/no-touch’, ‘running/no-touch’, ‘no-running/touch’ and ‘running/touch’) and categorized all neurons according to their highest response (see Experimental Procedures). 77.2 ± 9.6% of L2/3 neurons were integrative cells compared to only about half of the neurons in L5 (51.2 ± 4.5%). Accordingly, the two subpopulations of touch cells and run cells were larger in L5 than in L2/3 (Figure 6B-C). In an additional analysis we applied a general linear model to model the neuronal responses as the weighted sum of running-and stimulus-related parameters (see Experimental Procedures). Comparison of weights showed that L2/3 neurons gave larger values to both running and wall-touch parameters than L5 neurons (Figure S5). These results suggest that the neuronal network in L2/3 integrates locomotion and touch information to a higher degree compared to L5. At the population level, mean ΔF/F values of L2/3 neurons were similar for running/no-touch and no-running/touch conditions with an approximately linear summation for concomitant running/touch (Figure 6D). In comparison, mean ΔF/F values of L5 neurons were larger for running-only compared to touch-only condition, with only a marginal increase in population activity when wall-touching occurred during running. We conclude that L2/3 neurons show stronger integration of locomotion and ongoing touch signals than neurons in L5.

**Figure 6.**
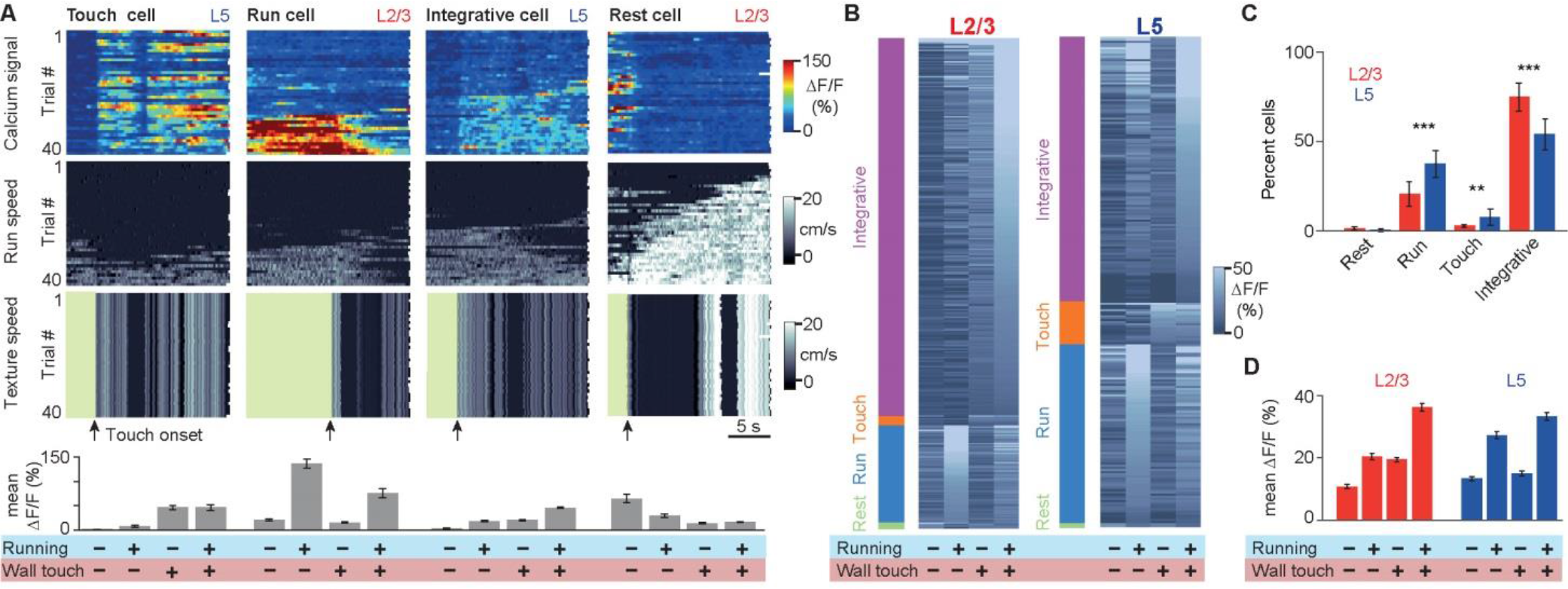
More L2/3 than L5 neurons integrate locomotion and concurrent wall touching. (A) Four example neurons with different response properties recorded during Open-loop stimulation. Each column presents data from a single neuron across trials in a single session. Top panel heat map shows ΔF/F calcium signal with each row representing a trial. Trials are sorted according to the mean run speed of a trial. Middle and bottom panels show the run speed and texture rotation speed of corresponding trials. Green periods in the bottom panel indicate when the texture was not in contact with whiskers. Bar graphs at the bottom are mean ΔF/F activity during the four stimulus and movement conditions: no-touch/no-running, running/no-touch, touch/no-running, and touch/running. (B) Categorization of all L2/3 (n = 338, 5 mice) and L5 (n = 236, 4 mice) neurons according to their activity during their first Open-loop session Color bar on the left show categories, gray scale shading shows mean ΔF/F at four stimulus/behavior conditions. Cells are sorted according to their mean activity during ‘running/wall touch’ condition relevant to their assigned category.

## DISCUSSION

Using a tactile virtual reality setup we found that neuronal activity across barrel cortex layers is strongly increased by running, more than by whisking. Furthermore, we found that continuous wall-touching evokes differential responses across layers, transient in L5 neurons and sustained in L2/3 neurons. In line with this finding the fraction of neurons best coding for running-with-touch was larger in L2/3 compared to L5. Moreover, in both L2/3 and L5 small subsets of neurons were sensitive to sensorimotor mismatches. Taken together, our findings highlight the strong influence of locomotion on barrel cortex activity and reveal a layer-dependence of touch responses in the active locomotion state.

### Running modulates barrel cortex activity more than whisking

Running strongly modulates sensory processing, but effects vary across sensory modalities and among cell types (see reviews by Busse et al., 2017; Händel and Schölvinck, 2017). We found that S1 neurons increase their activity during running (accompanied by whisking). In the absence of sensory stimulation, about 30% of both L2/3 and L5 neurons displayed a significant run-onset response, consistent with a previous study reporting similar effects mainly from L4 and L5 neurons (Sofroniew et al., 2015; the number of extracellularly recorded supragranular neurons were limited). Another study on somatosensory processing used running as a means to make mice continuously whisk against a stimulus bar but did not report on neuronal activity changes due to running (Pluta et al., 2015). In our experiments whisking minimally increased barrel cortex activity and we found only few whisking-onset responsive cells in both superficial and deep layers. Previous studies have reported inconsistent results regarding modulation of barrel cortex activity by whisking: on the one hand changes in membrane potential dynamics but not firing rate were observed (Crochet and Petersen, 2006; de Kock and Sakmann, 2009; but see O’Connor et al., 2010), on the other hand a prominent increase in firing rate upon whisking has been reported specifically for L5A neurons in rats (de Kock and Sakmann, 2009; O’Connor et al., 2010). Our findings are more consistent with the former results but further dissection of subtype-specific modulations in both superficial and deep layers is required.

Our results thus provide evidence that locomotion exerts strong effects on sensory processing in barrel cortex. One may therefore have to rethink the commonly applied notion that barrel cortex employs sparse coding (Brecht and Sakmann, 2002; Jadhav et al., 2009). The increased activity observed both in superficial and deep layers during running suggests that barrel cortex may employ state-dependent encoding strategies. In an ethologically relevant situation such as navigation, increased activity might provide more efficient and faster sensory coding.

### Layer-specific touch-evoked responses

Touch-evoked responses during running comprised initial strong activation of both superficial and deep pyramidal neurons but a subsequent decrease in activity with continuous stimulation only in deep pyramidal neurons. One explanation for such discrepancy in temporal response profiles between L2/3 and L5 could be intrinsic differences of neuronal classes. Indeed several studies reported differences in adaptation properties of subtypes of pyramidal cells. A recent study for example revealed distinct membrane depolarizations in S2- and M1-projecting L2/3 neurons: while S2-projecting neurons robustly signaled sensory information during repetitive active touch, M1-projecting neurons displayed strongly adapting postsynaptic potentials (Yamashita et al., 2013). In our experiments, a predominant activation of a larger population of S2-projecting neurons could explain the observed sustained response in L2/3. Similarly L5A neurons, which mainly send projections to other cortical areas, have an adapting spiking pattern whereas L5B neurons, whose projections target subcortical areas, have a regular spiking pattern upon current injection (Groh et al., 2010; Hattox et al., 2007; de Kock et al., 2007). The majority of our L5 imaging fields of views were localized around L5A. Thus, adapting responses of these cells might explain the observed transient activity upon continuous texture stimulation. On the other hand, electrophysiological recordings in rat barrel cortex while stimulating whiskers at different frequencies showed no significant difference in adaptation responses of superficial and deep layer neurons (Musall et al., 2014). This indicates that sustained versus transient responses may result not only from intrinsic differences of L2/3 and L5 neurons but in addition might involve circuit mechanisms on the local as well as long-range scale.

Inhibitory neurons within the local network might mediate differential response profiles across lamina. For instance, Pluta et al. (2015) reported that activation of L4 neurons, which in turn recruited a population of inhibitory fast spiking neurons in L5 (also see Gabernet et al., 2005), caused suppression of L5 neurons whereas activity of L2/3 neurons was increased. In this study mice were continuously running. Our results are in line with this observation and in addition suggest that recruitment of inhibition might be enhanced due to the running state of the mice because suppression was smaller when the animals were stationary. Another plausible mechanism to explain late suppression of L5 neurons is the recruitment of frequency-dependent disynaptic inhibition (FDDI) of Martinotti cells (Silberberg and Markram, 2007). In addition to L5 pyramidal cell inputs, L5 Martinotti cells receive strong input from L2/3 neurons (Jiang et al., 2015; Kapfer et al., 2007; Naka and Adesnik, 2016). Considering more sustained activity of L2/3 neurons upon continuous wall-touch, interlaminar activation of L5 Martinotti cells might be more significant. Additionally this inhibition process is weak at the beginning but is facilitated with continuous input (Tan et al., 2008), thereby possibly accounting for the delayed suppression of L5 pyramidal neurons. Several studies have also suggested that locomotion effects are mediated via disinhibition of pyramidal neurons by activation of vasoactive intestinal peptide (VIP) expressing interneurons upon locomotion (Fu et al., 2014; Pakan et al., 2016). VIP neurons are more numerous in superficial layers, albeit their axons can reach different layers. Hence, their activation might lead to more effective disinhibition in L2/3 upon running compared to L5, creating a disparity in responses of superficial and deep layer neurons.

Finally, distinct long-range afferent inputs to L2/3 versus L5 neurons in general could underlie the distinct response profiles (Aronoff et al., 2010; Kinnischtzke et al., 2014; Mao et al., 2011; Petreanu et al., 2009). For instance POM sends axons to L1 and L5 (Lu and Lin, 1993; Wimmer et al., 2010). Although the specificity of targeting of L1 axons onto apical dendrites of both L2/3 and L5 neurons remains unclear, specific inputs to L5A could explain distinct response patterns. To our best knowledge changes to POM activity upon running remain to be investigated. In addition layer-specific inputs originating from motor cortex to somatosensory cortex may contribute to response diversity.

### Functional sensorimotor representations across layers

Laminar organization of a cortical column in the mammalian neocortex is highly preserved throughout evolution and it is of key importance to compare response properties across layers to better understand their functional role. One suggested role of laminar organization in cortex is that it may provide a framework for hierarchical processing such that a ‘higher’ cortical area sends expectation information through axonal projections to superficial neurons of a ‘lower’ sensory area, where top-down inputs are integrated or compared with bottom-up inputs (Bastos et al., 2012). According to this model, one might expect expectation mismatch signals to be more prominent in superficial than in deep layers. In this study we indeed found evidence of mismatch responses in barrel cortex but we did not observe differences across lamina. In our experiments around 5% of both superficial and deep layer neurons in S1 showed increased activity to a mismatch between the animal’s running speed and texture speed. Abrupt stalling of tactile flow while the animal was stationary did not cause increased activity. An equivalent study in primary visual cortex reported that a similar subset (13%) of L2/3 neurons respond to expectation mismatch in visual stimuli, a brief stalling of the visual stimulus flow while the animal is running on a treadmill (Keller et al., 2012). In visual cortex, to our best knowledge, there is no comparison of mismatch responses of L2/3 neurons with that of L5 neurons. In visual cortex, mismatch responses were observed also at population level, different from in our experimental results showing a decrease in population activity upon mismatch, which can be explained by a larger fraction of neurons decreasing their activity upon texture rotation stall.

Within the local microcircuitry of barrel cortex we found a distributed representation of sensory information (textured wall) and motor information (running). The majority of neurons, especially in L2/3, responded strongest for the combined condition of run-with-touch. In L5 half of the neurons responded best to a single modality, wall-touch or running. This finding is also consistent with the running-onset responses as well as the modulations of mean ΔF/F values, with L5 neurons showing larger increases upon running. Additionally, fewer integrative cells in L5 are in line with the suppression of L5 neuron activity with continuous touch. Overall these findings provide direct evidence for a more integrative role of superficial neurons during sensory processing in S1 and suggest that L2/3 neurons play the primary role during contextual modulation of cortical activity and multimodal integration of sensory inputs.

### Future perspectives

Overall our findings suggest that L2/3 neurons enter a continuous monitoring mode during running while L5 neurons mainly respond to salient changes. We therefore speculate that L2/3 and L5 neurons might contribute differentially to behavioral tasks with different requirements. For example, L5 neurons might be more involved in obstacle detection, requiring immediate changes in the motor program, while activity of L2/3 neurons might be critical for texture-discrimination during navigation. In future experiments, it will be interesting to verify distinct task involvement of L2/3 and L5 populations. In addition, the contributions of different subtypes of pyramidal cells as well as inhibitory interneurons to shaping the response dynamics of cortical layers during sensorimotor integration warrant further investigation.

## ACKNOWLEDGEMENTS

We thank Hansjörg Kasper, Martin Wieckhorst, and Stefan Giger for technical assistance and Yaroslav Sych, Ariel Gilad and Christopher Lewis for comments on the manuscript. This work was supported by grants from the Swiss National Science Foundation (SNSF) (31003A-149858 and Sinergia grant CRSII3_147660; F.H.), the European Research Council (ERC Advanced Grant BRAINCOMPATH, project 670757; F.H.), Sir Henry Dale Fellowship jointly funded by the Wellcome Trust and the Royal Society (Grant 200501/Z/16/Z; A.B.S.), SNSF Marie Heim-Vögtlin grant (PMPDP3_145476; A.A.), SNSF Ambizione grant (PZ00P3_161544; A.A.).

## AUTHOR CONTRIBUTIONS

Conceptualization, A.A. and F.H.; Methodology, A.A; Investigation, A.A., and A.S.; Software and Formal Analysis, A.A.; Writing – Original Draft, A.A. and F.H.; Writing-Review & Editing, A.B.S., A.S., A.A. and F.H.; Funding Acquisition, A.A.; Resources, A.A. and F.H.; Supervision, A.B.S, A.A., and F.H.

## EXPERIMENTAL PROCEDURES

All experimental procedures were conducted in accordance with the ethical principles and guidelines for animal experiments of the Veterinary Office of Switzerland and were approved by the Cantonal Veterinary Office in Zurich.

### Virus injections and cranial window preparation

We used 5-9 weeks old male C57BL6 mice. Virus injection and cranial window preparation followed a former description (Chen et al., 2013) and were performed under isoflurane anesthesia (1.5-2%) with body temperature maintained at approximately 37°C using a regulated heating blanket and a thermal probe. The eyes of the mouse were covered by Vitamin A cream (Bausch & Lomb) during the surgery. After hair removal and disinfection with ethanol (Alkopads B.Braun), the skin was opened with a scalpel and the exposed cranial bone was cleaned from connective tissue and dried with cotton pads (Sugi). To express the red calcium indicator R-CaMP1.07 (Ohkura et al., 2012) in cortical neurons, we injected AAV2.1-EF1α-R-CaMP1.07 into barrel cortex (at 3.3 mm lateral and 1.1 mm posterior to bregma). Two injections of 210 nl of viruses (approximately 1.21 x 1013vg/ml) were performed across 300-700 µm below the pial surface to achieve expression in both deep and superficial cortical neurons. Afterwards a circular piece of cranial bone (Ø 4 or 5 mm) was removed using a dental drill, leaving the injection sites in the center. A coverglass (Ø 4 or 5 mm, 0.17 mm thickness) was inserted and secured in place by UV curable dental acrylic cement (Tetric Evoflow). In order to ensure reproducible positioning of the mouse by head-fixation under the microscope objective, a small aluminum hook was glued to the skull on the contralateral side of the head with dental cement. Intrinsic optic signal imaging was used to verify viral expression area within barrel cortex (Chen et al., 2013). The barrel field of single whiskers (mainly B1, B2, C1 and C2) for each mouse was identified under light anesthesia (approximately 0.5-1% isoflurane).

### Tactile virtual reality setup

Rodents are highly tactile animals. In their natural environment they run through dark tunnels utilizing their whiskers to touch the walls. To simulate this natural behavior, we built a tactile virtual reality setup (Thurley and Ayaz, 2017). Mice were head-restrained on a rung ladder treadmill (Ø 23 cm) with regularly spaced rungs (1 cm spacing). Run speed and distance were recorded at 40 Hz with a rotary encoder (incremental 5 VDC 360, RI32-O/360AR.11KB, Hengstler). Textures (sandpapers of various graininess: P100 or P1200) were presented on rotating cylinders and were brought in reach of the whiskers with a linear motorized stage (Zaber T-LSM050B stage with built-in controller). Texture contact was established after mice had run a predefined distance on the treadmill. This distance (5 – 50 cm) was determined for each mouse individually, to allow several seconds of recording before texture touch. A stepper motor (Phidgets 3305_5 NEMA-17 Bipolar 20 mm Stepper) was used to control the speed of the texture rotation. The texture speed was either coupled (Closed-loop) or de-coupled (Open-loop) to the animal’s run speed. Whiskers on the contralateral side of the cranial window were illuminated with 850-nm infrared LED light while being monitored with a CMOS high-speed camera (Optronis, CL600X2) at 200-Hz frame rate. The behavioral set up was controlled by custom software developed in LabVIEW. This software served as the master control unit for controlling and recording behavioral components and triggered whisker monitoring and two-photon calcium imaging.

### Behavioral paradigms and experimental design

Animals were free to move on the treadmill and did not receive any reward under any condition. After habituation whiskers were trimmed such that they would not contact the treadmill to avoid somatosensory stimulation by the ladder rungs. We considered three behavioral and stimulation conditions (see Results section in main text for details of conditions): (1) ‘No-touch’ (5 mice, 10 imaging spots; up to 3 sessions per spot) (2) ‘Closed-loop’ (5 mice, 12 imaging spots; up to 9 sessions per spot) and (3) ‘Open-loop’ (5 animals, 11 imaging spots; up to 4 sessions per spot). To control for confounding effects of the sounds due to motor rotation and linear-stage movement, we rotated the textured cylinder and moved the linear stage in ‘No touch’ trials as if in ‘Closed-loop’ condition but kept the textured ‘wall’ out of reach for the whiskers.

A week after the cranial surgery, mice were first habituated to the experimenter by handling. Once familiar, animals were accustomed to the behavioral set up, where they freely moved on the rung ladder treadmill while being head-fixed and presented with Closed-loop rotating texture stimuli. After a week of habituation and training we performed two-photon calcium imaging of neuronal populations in superficial and deep layers of S1 barrel cortex (221-664 µm below the pial surface) during several sessions under all three behavioral conditions. Each behavioral session was composed of 20-s long trials with 3-5 s inter-trial intervals. The number of trials in each session varied from 10 to 80. Animals were kept under these recording conditions maximally for 45 min per recording session and about 1.5 hours per day in multiple sessions. For each imaging area, we collected data in 1-6 experimental sessions spread over maximally 10 days. For repeated imaging across several days, individual cells were reidentified by their shape and localization relative to the spatial constellation of the cells in the neighborhood within the imaging field.

### Two-photon calcium imaging

We used a custom-built two-photon microscope of the Sutter Movable Objective Microscope (MOM) type. This system was equipped with galvanometric scan mirrors (model 6210; Cambridge Technology) and a Pockels cell (model 350/80 with controller model 302RM, Conoptics) controlled by HelioScan software (Langer et al., 2013). The objective was a water immersion 16× objective (CFI LWD 16×/0.80; Nikon). For excitation of R-CaMP1.07 we used a ytterbium-doped potassium gadolinium tungstate (Yb:KGW) laser (1040 nm; 2.5-W average power; ~230-fs pulses at 80 MHz; model Ybix; Time-Bandwidth Products). Fluorescence was collected through a red emission filter (610/75 nm; AHF Analysentechnik) and detected by a GaAsP PMT (Hamamatsu H10771P-40 SEL). We performed in vivo calcium imaging using 33-160 mW average power below the objective for focal depths ranging from 221-664 µm. Image acquisition rate was 10 Hz over a 140-µm by 180-µm field of view.

### Slice histology and confocal microscopy

After the last in vivo experiments mice were anesthetized by i.p. injection of ketamine (0.15 mL, 50 mg/mL). 0.05 mL heparin was injected in the left hearth ventricle and the animal were intracardially perfused with 20-25 ml of phosphate buffer (0.1 M, pH 7.3, room temperature) and subsequently with 20-30 ml of paraformaldehyde solution (PFA; 4% in 0.1 M PBS, pH *7.3,* room temperature) both at 11 ml/min. The brain was extracted and postfixed in 4% PFA at 4°C overnight. Afterwards, it was rinsed three times with phosphate buffer and preserved in 30% sucrose (0.1 M phosphate buffer) at -20°C until further processing. After unfreezing the brain was cut into 50-µm free floating coronal slices with a microtome (Leica VT1000 S). Brain slices were mounted on microscope slides, embedded in Fluoromount (Dako), and covered by a glass cover. We acquired fluorescence stacks (3-µm z-steps) of R-CaMP1.07 expression with a confocal laser-scanning microscope (Olympus FV1000; 546-nm excitation wavelength).

### Data analysis

#### Calcium imaging data analysis

Frames in the time series of two-photon imaging data were registered using a Hidden-Markov-Model, line-by-line motion-correction algorithm (Dombeck et al., 2007). Regions of interests (ROIs) corresponding to individual neurons were manually selected from the mean image of a single-trial using Fiji software (Schindelin et al., 2012). Image frames of other trials were then realigned to the reference trial, from which the ROIs were selected, to account for possible shifts of the field-of-view throughout an imaging session. R-CaMP1.07 fluorescence signals and behavioral data were analyzed using custom MATLAB scripts (Mathworks). Background fluorescence (estimated as bottom 1st percentile of the fluorescence signal across the entire movie) was subtracted from all trials. Insufficiently motion-corrected trials or time periods within a trial were excluded from the analysis. In rare cases, ROIs with large motion artifacts were also excluded. Background-and motion-corrected R-CaMP1.07 signals were expressed as relative percentage change of the fluorescence ΔF/F=(F–F0)/F0, where baseline F0 was calculated as the 1st percentile of the smoothed fluorescence trace (51-point 1st-order Savitsky-Golay filter) after concatenating fluorescence signals over all trials within a session. Smoothing over a large window aimed to estimate the baseline fluorescence of a neuron when it was inactive, likely not firing action potentials. Similarly we defined the baseline noise, σ, as the standard deviation of the fluorescence change during the least noisy 5-s period within a session (1st percentile). We used σ for identifying neurons responsive to salient events (e.g. whisking onset, running onset, touch or perturbation).

### Whisking and running analysis

We monitored whisker motion at 200 Hz and measured the mean whisker angle across all imaged whiskers using automated whisker tracking software (Knutsen et al., 2005). Whisker angle traces were down-sampled to the two-photon imaging frame rate of 10 Hz. We assigned whisking and no-whisking periods based on whether the standard deviation of the mean whisker angle trace (calculated in a 51-point sliding window) was above or below a predefined threshold (>2.5º). An encoder recorded run speed at 40 Hz and the run speed trace was smoothed with a 1-s Gaussian filter and also down-sampled to 10 Hz. To assign running versus resting periods, run speed traces were further smoothed with a 1.1-s broad 1st-order Savitsky-Golay filter and periods with absolute values of run speed >0.8 cm/s were considered as ‘running periods’.

We analyzed modulatory effects of whisking and running on barrel cortex activity under ‘No-touch’ condition. To determine the effect of whisking alone we excluded any running period. We defined the whisking modulation index (MI) as

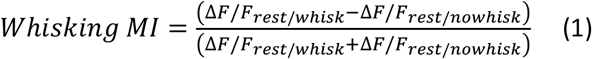

Where Δ*F/F_rest/whisk_* and Δ*F/F_rest/no whisk_* denote the mean ΔF/F value during corresponding resting/whisking and resting/no-whisking states, respectively. Similarly, the effect of running was assessed by comparing mean ΔF/F values during running versus resting states independent of the whisking state of the animal. Running MI was calculated analogous to whisking MI as

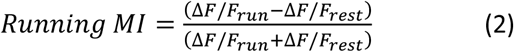

Where Δ*F/F_run_* and Δ*F/F_rest_* denote the mean ΔF/F values in the respective time windows.

### Event detection and signal alignment

For detection of the whisking and running onsets we used the binary running/resting and whisking/no-whisking vectors. Any at least 400-ms long period of no-whisking (or resting) period followed by at least 400-ms of whisking (or running) was detected as a whisking onset (or running onset, respectively). To capture the time course of whisking-evoked ΔF/F responses independent of running we considered onset-aligned ΔF/F traces from 2 s before whisking onset until either whisking stopped or the animal started running. Similarly running-evoked ΔF/F responses were considered from 2 s before running onset until the animal stopped running. As animals were almost always whisking when they were running, it was not possible to isolate running from whisking. Because of uncertainties in determining the first actual texture-whisker touch moment, we defined touch onsets as the time points where the linear stage carrying the texture started to move towards the whiskers. The intact set of whiskers together with whisker motion prevented us from detecting the precise moment of first touch of the whisker corresponding to the imaged barrel column. Our rough estimate of the actual time of first touch (of any whisker) is at 158 ± 48 ms after the texture cylinder started moving in (reaching its final position to 245 ± 44 ms; mean ± s.d.). Finally, perturbation events occurred at the beginning of the 2-s window when texture rotation was halted. To compute the perturbation response we consider only events that occurred when the mean run speed of the animal was larger than 2 cm/s. This choice of threshold ensured that a mismatch between run-speed and tactile flow was indeed imposed by the perturbation.

We averaged event-aligned ΔF/F traces across trials (smoothed with 1st order Savitsky-Golay filter, 500-ms window) and calculated the event modulation as the difference in mean ΔF/F values between pre-and post-event windows. For whisking, running and touch modulations we calculated the difference in as the max ΔF/F in the post-event window minus the pre-event mean ΔF/F. Perturbation modulation was computed from the mean ΔF/F values in both windows. Neurons with modulations larger than twice the baseline noise (2σ) were considered responsive. Specific calculations for each event type are as follows:

For Open-loop condition touch events were divided into ‘touch during running’ and ‘touch during resting’ groups. For each event we computed the mean running speed within a 5-s window (−1 to +4 s) around the touch onset. Events with mean speed > 2 cm/s were selected as ‘touch during running’ events and ones < 1 cm/s was considered ‘touch during resting’.

If a neuron was imaged over several sessions we randomly selected a session and considered each neuron once in any population analysis. Averaging event triggered responses over several sessions gave qualitatively gave similar results (data not shown). Only for perturbation responses we selected the session the neurons was most responsive. Hence our perturbation results are an over-estimation.

### Functional classification of neurons

For functional classification of neurons we considered only Open-loop sessions as they comprised various combinations of stimulation and locomotion behavior. Calcium signals were smoothed with a 1st-order Savitsky-Golay filter with 500-ms window and resampled 100 times by selecting equal number of trials of the original session by replacement. For each selection of trials mean ΔF/F and standard deviation were calculated for 4 different stimulus-behavior conditions: no-running/no-touch; running in the absence of wall-touch; wall-touch without running; and concurrent running and wall-touch. Cells which show the largest activity in any of these categories was labeled as stationary, run, touch and integrative cells, respectively. Cells were initially assigned to the category where their mean ΔF/F was the largest. But if the difference in mean ΔF/F between first and second largest response was smaller than one standard deviation of the largest response category cell was assigned to the second largest group. This aimed to assign cells to single component category (run or texture touch) unless their response significantly increased with the addition of the second component. Final classification of cells, for the first time they were imaged, is shown in Figure 6B (categories computed with concatenated multiple Open-loop sessions gave equivalent distributions, data not shown).

This analysis involved 4 animals, 24 sessions, 11 different imaging areas of 574 cells (338 L3 and 236 L5). 2 sessions were not included where 1 of these 4 categories were not realized (i.e. there were no periods of texture touch without running). Statistics of cell category distribution was performed by randomly selecting 11 of 24 sessions 10 times. For each selection percent cells in each category was calculated separately for L2/3 or L5 populations. 10 selections provide mean and standard deviation of percent cell category distributions. For each category, two sampled t-test was performed between percent representations in L2/3 and L5 populations.

### General linear model of neuronal responses

We tried to capture behavior and stimulus related activity of neurons using a general linear model (GLM). We considered Open-loop experiments in our modeling approach as this condition brought together different combinations of run speed, wall contact, and texture rotation speed. We expressed neural calcium activity for each neuron as the weighted sum of these variables:

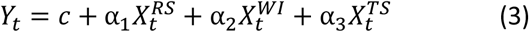

Here we used the run-speed of the animal 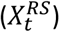 the binary vector representing whether the wall is in reach or not 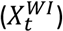, and the texture rotation speed 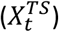 to model the smoothed (1 second moving average) calcium signal *(Y_t_)* of each neuron at time t.

We considered 24 Open-loop sessions from 5 mice and some neurons are represented multiple times if they were recorded from in multiple Open-loop sessions. Model was fit using 90% of the data (training set) and tested on the remaining 10% (test set). This process was repeated 10 times for each neuron. The quality of the fit was computed for each selection (i) of data sets as the fraction of variance explained (Qi):

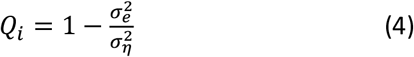

where σ_e^2 is the mean squared error of the model predictions for the test set and σ_η^2 is the variance of the training set. They are defined as:

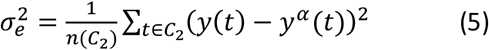

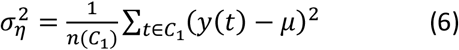

where y(t) is the smoothed calcium signal of the neuron at time t, *y^α^(t)* is the prediction by the model given parameters Xt and coefficients αi, C1 and C2 are training and test sets, respectively. Here μ is the mean firing rate of the training data. A higher Q suggests better performance. Values of Q close to 1 are unlikely due to the intrinsic variability in the response of neurons, which is uncorrelated to the stimulus or behavior. Very low values of Q suggest that the model is unreliable. Therefore, we only considered neurons whose responses were predicted with σ > 0.15, where σ is the mean of all Q. 453 of 626 L2/3 neurons and 248 of 416 L5 neurons satisfied this criterion.

## Supplementary Figures

**Figure S1.**
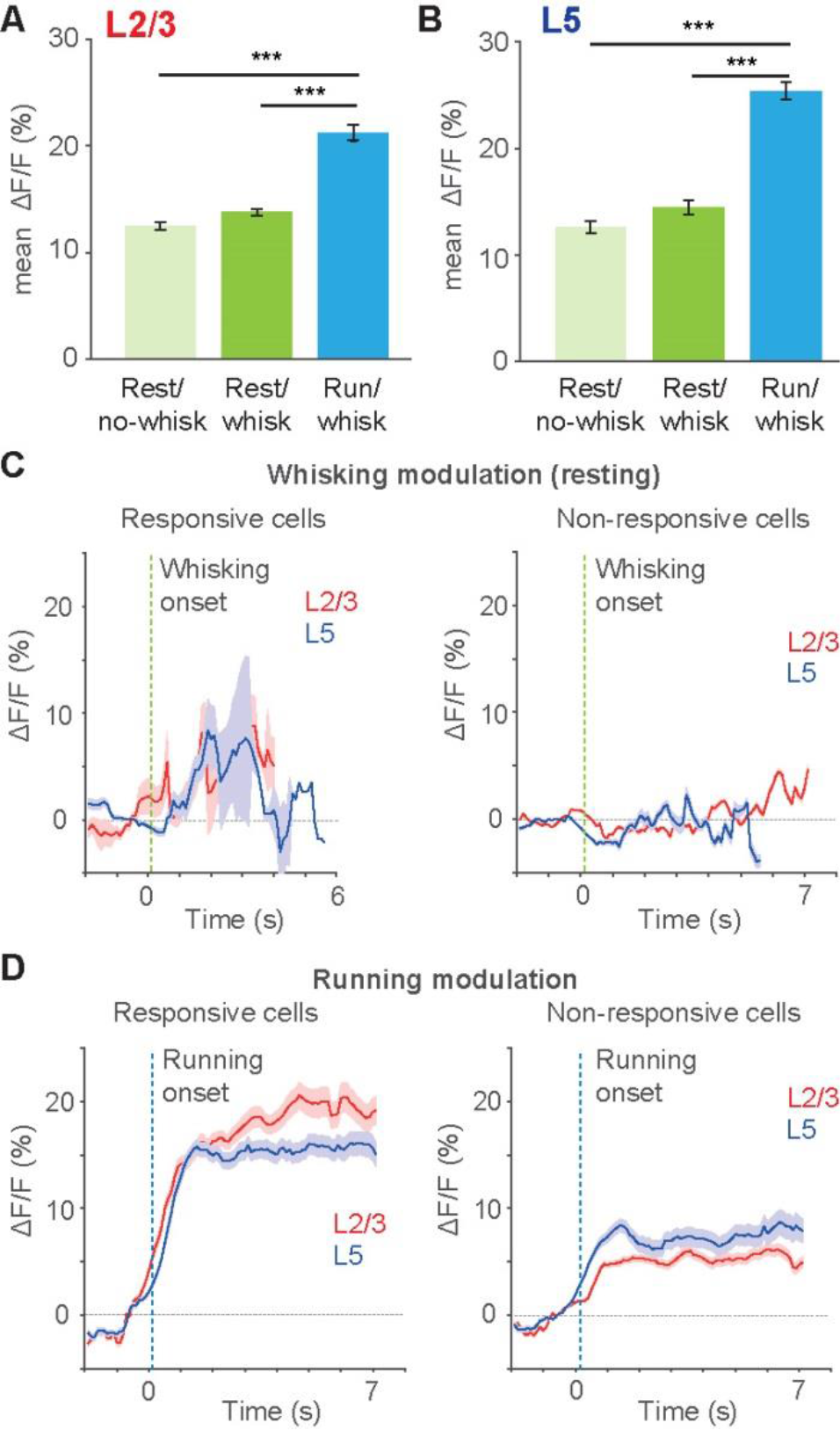
Running increases S1 activity. (A) Comparison of mean ΔF/F for L2/3 neurons during resting/no-whisking, resting/whisking and running/whisking periods in the absence of texture stimulus. For L2/3 neurons mean ΔF/F values during corresponding behavioral conditions were 12.5 ± 0.4%, 13.7 ± 0.4% and 20.8 ± 0.7%, respectively (±s.e.m., n = 342 neurons). Statistical significance is computed with multi-sample comparison with one-way ANOVA. *** p < 0.01.
(B) Same as in A but for L5 neurons. Mean ΔF/F values were 12.7 ± 0.6%, 14.5 ± 0.7% and 25.4 ± 0.9% for corresponding behavioral states for L5 neurons (n= 168).
(C) Comparison of whisk onset responses of responsive neurons vs non-responsive neurons, where responsive neurons elicit onset modulation larger than 2σ (σ is the baseline noise; see Experimental Procedures; here 7.6 ± 1.7% for L2/3 and 8.3 ± 2.0% for L5 neurons, respectively; mean ± s.d.).
(D) Similar to C but comparing run onset responses for responsive and non-responsive neurons. Increased activity upon run onset in non-responsive neurons indicates that larger population show increased run onset modulation but the extent of increase was not large enough for our selection criteria (> 2 σ), which was quite conservative

**Figure S2.**
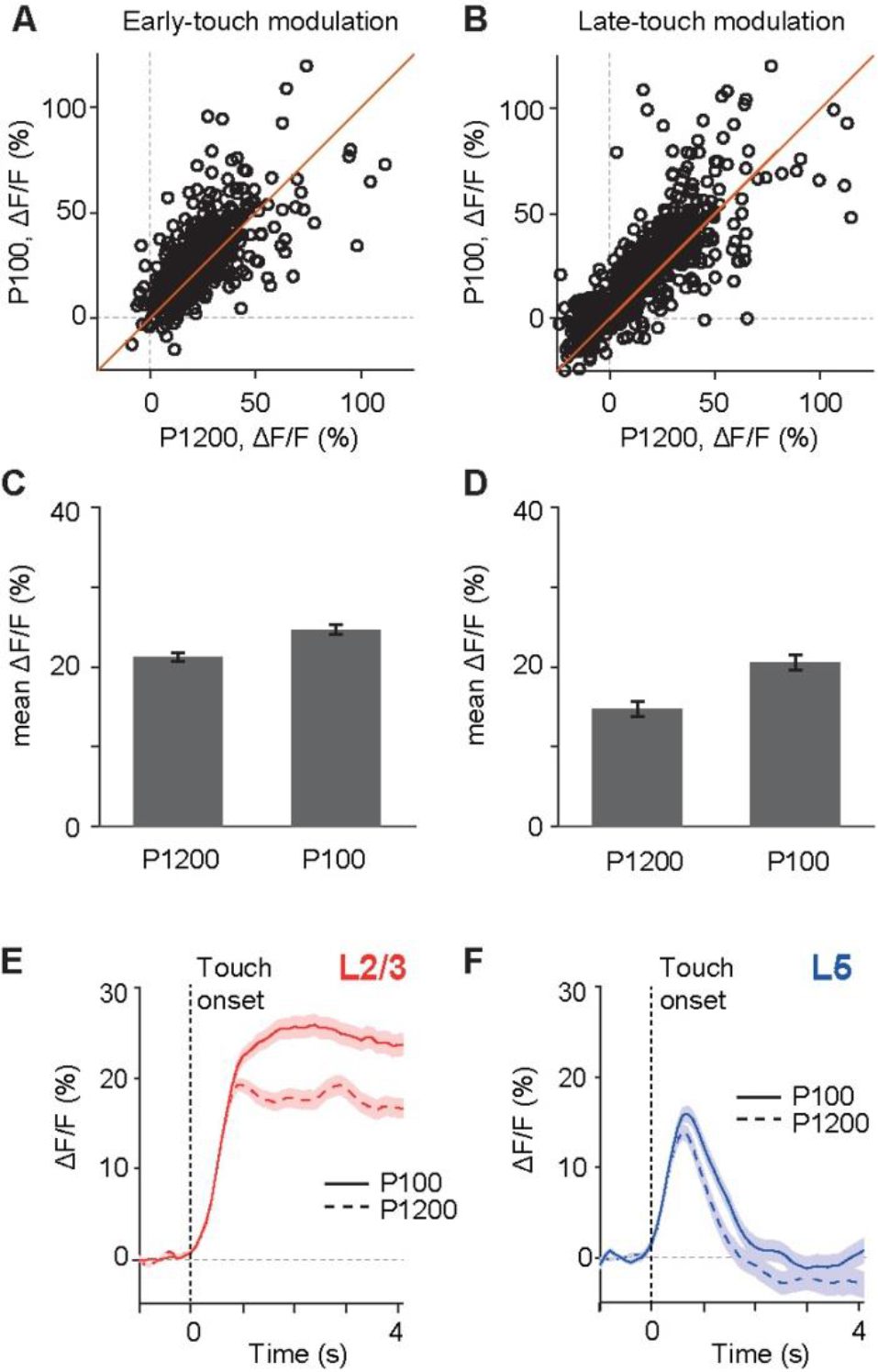
Wall-touch responses to different textures. We used sandpapers of two different graininess (P100-rough and P1200-smooth) in our touch experiments. Sessions either involved a single texture or random representations of both textures in each trial. Here we compare neuronal responses to wall touch of random presentation of two different textures (14 Closed-loop sessions, n = 691, neurons might be repeated). (A) Scatter plot of early-touch modulations of wall touch events where the wall was textured with P1200 (x-axis) and P100 (y-axis) sandpaper.
(B) Same as in A but comparing late-touch modulations.
(C) Population averages of early-touch modulations elicited by smooth and rough textures, respectively (21.4 ± 0.5% ΔF/F vs. 24.9 ± 0.6% ΔF/F, mean ± s.e.m., p < 10^−4^ paired T-test).
(D) Same as in C but for late-touch modulations (15.0 ± 0.9% ΔF/F for P1200 vs. 20.9 ± 1.0% ΔF/F for P100, mean ± s.e.m., p < 10^−4^ paired T-test).
(E) and (F) Comparison of population averages of touch responses for rough (P100) and smooth texture (P1200) for L2/3 (n = 465) and L5 (n = 226) neurons, respectively. Although the rough texture elicited a slightly larger responses, consistent with previous findings (Chen et al., 2015), the difference in sustained versus transient touch responses of L2/3 and L5 neurons was not affected by texture identity.

**Figure S3.**
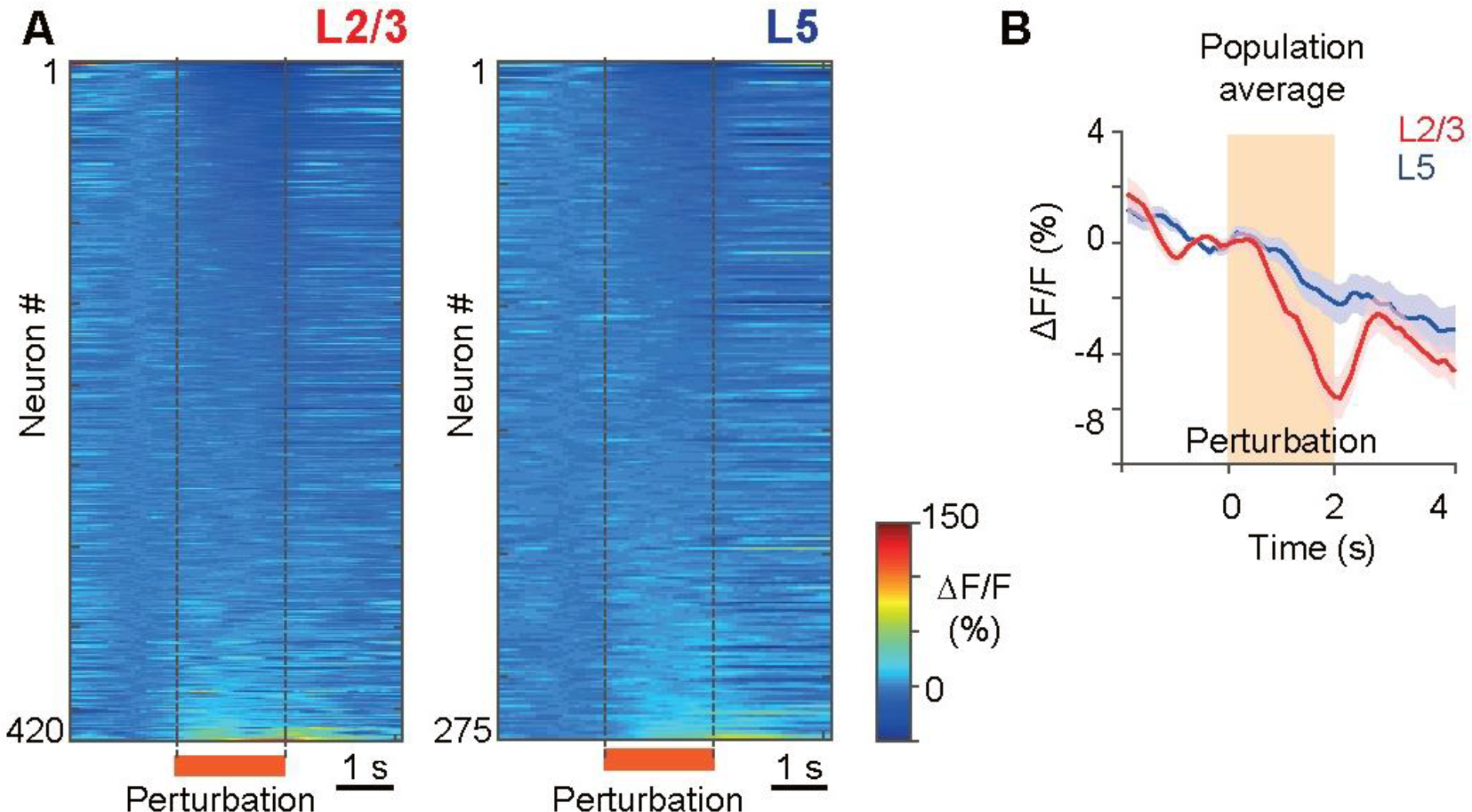
Decrease in activity dominates population responses to mismatch of running speed and tactile flow. (A) The heat map represents perturbation-evoked responses of all L2/3 (left column) and L5 neurons (right column) in Closed-loop condition. Each row represents the fluorescence change of a single neuron compared to the mean activity during a pre-perturbation window (−1 to -0.3 s). Orange period is when tactile flow was stalled suddenly.
(B) Population average of all neurons aligned at perturbation onset.

**Figure S4.**
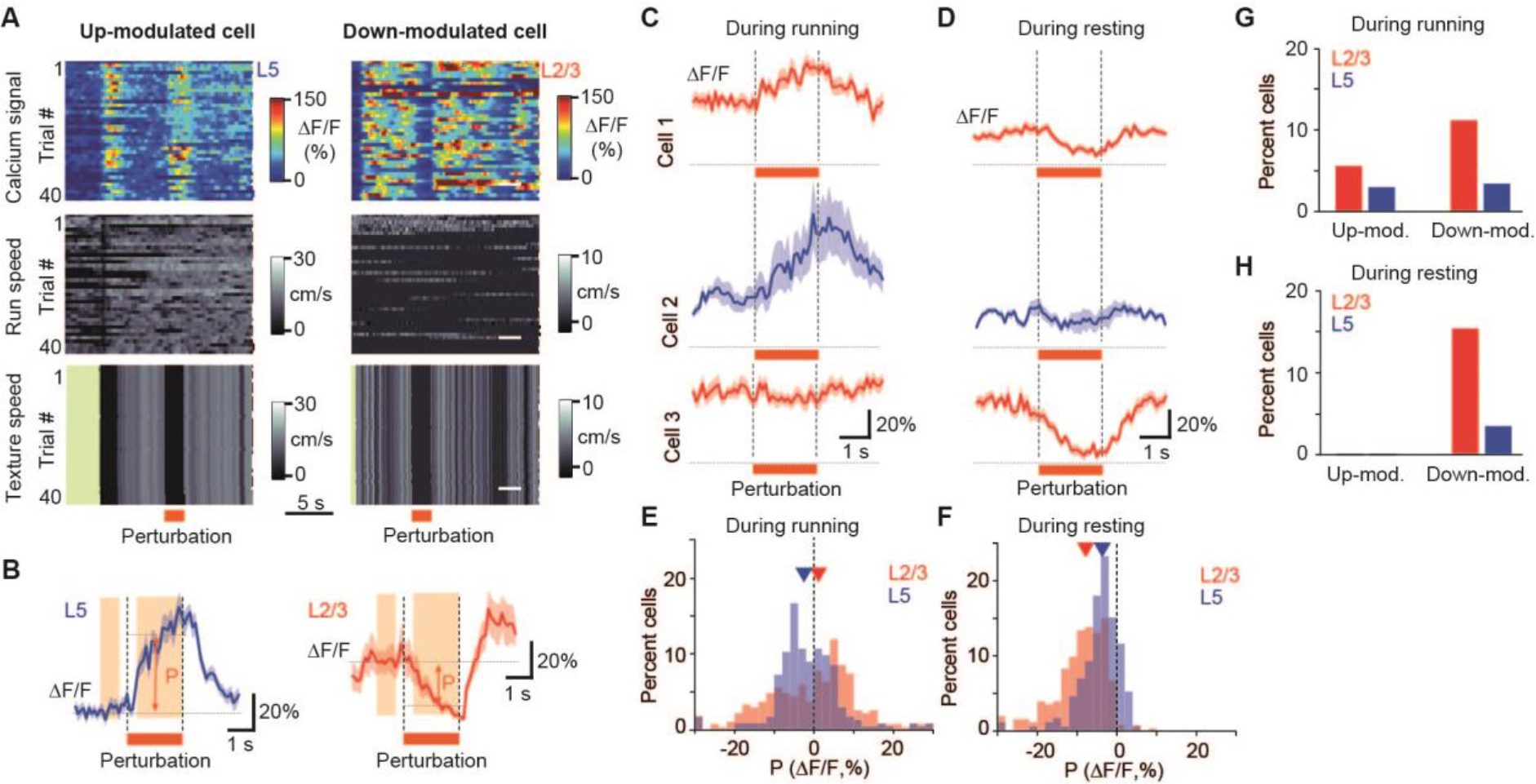
Perturbation-evoked responses during Open-loop condition. (A) Example responses of an up-modulated (left column) and a down-modulated (right column) neuron upon replay of a trial including a sudden stall of texture rotation (‘perturbation’) in Open-loop condition. Heat maps show calcium activity along with the running speed and the texture rotation speed over 40 trials.
(B) Average activity aligned at perturbation onset are plotted for example neurons in A (mean ± s.e.m.). P indicates the perturbation modulation defined as the difference between mean ΔF/F before and during perturbation (orange shaded windows) where running speed was larger than 2 cm/s.
(C) and (D) In Open-loop, sudden stalling of texture rotation may happen during running or resting. Here we present responses to ‘texture rotation stall’ during running (C) or during resting (D) of 3 example cells.
(E) Histograms indicate perturbation modulation (P) distributions for L2/3 (red, n = 269) and L5 (blue, n = 234) neurons during running (mean P equals -0.7 ± 0.7% and -1.7 ± 0.6% for L2/3 and L5, p < 0.05).
(F) Same as in E but for resting state P modulations for L2/3 (red, n = 253) and L5 (blue, n = 172) neurons during running (mean P equals -9.2 ± 0.5% and -4.3 ± 0.4% for L2/3 and L5 respectively, p < 0.0001).
(G) Percentage of neurons that are significantly up-modulated (P > 2σ) and down-modulated (P < -2σ) for L2/3 (red) and L5 (blue) populations during running.
(H) Same as in G but during resting state.

**Figure S5.**
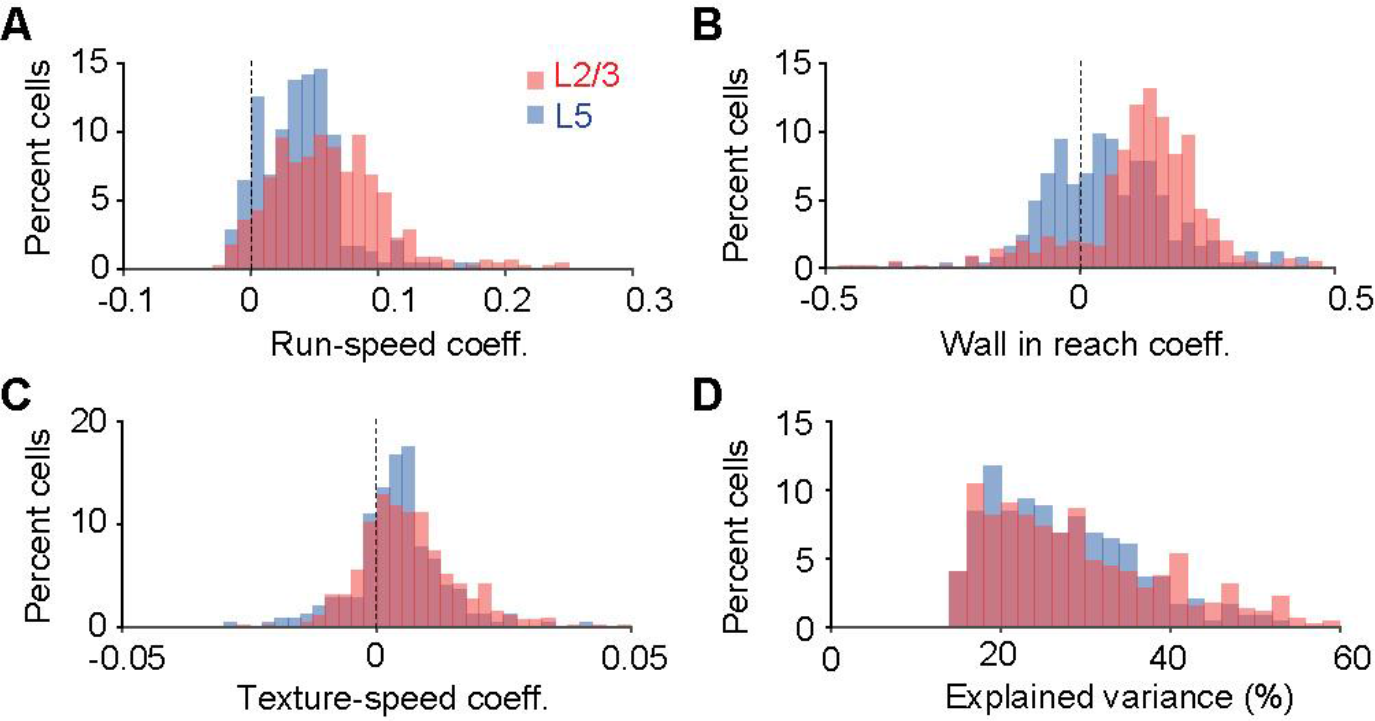
General linear model (GLM) parameter comparison supports more integrative role for L2/3 neurons. We modeled calcium signals from each neuron acquired during Open-loop sessions as a weighted sum of run-speed of the animal, the binary vector representing whether the wall is in reach of whiskers, and the texture rotation speed. We considered 24 Open-loop sessions from 5 mice and some neurons are represented multiple times if they were recorded from in multiple Open-loop sessions. Quality of the fit was quantified as the mean explained variance (Q) of 10 cross-validation data sets. This figure presents results from 453 of 626 L2/3 neurons and 248 of 416 L5 neurons with explained variance larger than 0.15 (neurons might be repeated). (A), (B) and (C) compare distributions of run-speed, wall-in-reach and texture-speed coefficients for L2/3 (re) and L5 (blue) populations. (D) Distribution of explained variance of model fits (Q > 0. 15).

